# Anthropogenic disturbance and spatial heterogeneity shape vegetation diversity in tropical swidden mosaics globally

**DOI:** 10.1101/2025.03.05.641554

**Authors:** Shane A. Scaggs, Xinyi Wu, Zaarah Syed, Denis Tverskoi, Jensan Lebowitz, Rongjun Qin, Sean S. Downey

## Abstract

Swidden agriculture is a widespread anthropogenic disturbance regime in tropical forests. Swidden research often posits that aggregate levels of forest disturbance correlate with increased land degradation, however a landscape configuration approach may distinguish when swidden degrades landscapes and when it diversifies them. Here we analyze how the configuration of swidden mosaics relates to vegetation diversity. Using satellite imagery from 18 swidden societies across the African, Southeast Asian, and American tropics, we quantify patch geometry using landscape metrics and estimate vegetation diversity from spectral variation and develop a nonlinear hierarchical Bayesian model that links the structure of swidden mosaics with vegetation diversity. Our analyses reveal three dominant gradients of swidden mosaic patterns: (1) aggregation versus interspersion of land-cover types; (2) spatial dispersion versus synchronization of disturbed patches; and (3) alternative modes of landscape connectivity. Across sites, vegetation diversity exhibits a consistent nonlinear response, peaking at intermediate levels of disturbance intensity. These results demonstrate that swidden agriculture does not produce a singular, degradative outcome. Instead, its effects on vegetation diversity depend on how disturbance is spatially configured. By shifting attention from area-based measures of deforestation to landscape configuration, this study reframes swidden as a spatial process with the potential for diversity-enhancing outcomes.

## 1 Introduction

Swidden agriculture, a rotational system of forest clearing, cultivation, and vegetation regrowth, creates visually distinctive spatial mosaics that can be understood from the perspectives of landscape and disturbance ecology, as swidden involves repeated biomass clearing events that interact with ecological recovery and successional processes to generate spatial heterogeneity. While local case studies have documented swidden with considerable descriptive detail [1], and remote sensing studies have revealed the extent of swidden activities globally [2, 3], few studies have examined how spatial heterogeneity varies systematically across swidden systems (although [4] is a noteworthy exception). Instead, studies of swidden landscapes have focused on aggregate levels of disturbance while ignoring the the complex and dynamic nature of this land use practice [5, 6]. By looking at swidden mosaics through the lens of landscape ecology, we adopt concepts and tools for understanding spatial patterns of land use and vegetation [7] that shift the focus toward the spatial configurations that arise from this dynamic agricultural technique. In this paper, we apply these ideas by quantifying spatial configuration and vegetation diversity in an attempt to understand how anthropogenic disturbances shape spatial heterogeneity in swidden mosaic landscapes.

Understanding how swidden agriculture shapes spatial heterogeneity is critical for supporting the conservation of biodiversity [8], the livelihoods of remote, rural, and Indigenous peoples [9] and responding to the impacts of global climate change [10]. Because agricultural systems which generate spatially complex landscapes have the potential to support significant plant and animal diversity [11, 12], we might expect similar potentials in swidden mosaics. Ethnobotanical studies have documented substantial plant diversity within swidden landscapes [13, 14] that is not limited to cultivated species [15]. Historical swidden disturbances appear to contribute to the formation of vegetation mosaics with increased heterogeneity [13, 16]. Such disturbance-driven mosaics have the potential to create novel ecotones and gradients that facilitate dispersal, foraging, and habitat availability [17, 18]. When a portion of mature forest is maintained, these matrices of mature and successional vegetation may also serve as essential carbon storage sites [19, 20]. Here we deepen our understanding of the ecological role of swidden agriculture by conceptualizing it as a widespread example of an anthropogenic disturbance process and studying its influence on vegetation diversity and spatial heterogeneity.

To this end, we focus on two primary research questions in this study:

1. What are the structural characteristics of swidden landscape mosaics and how do they vary across tropical regions?
2. How are local patterns of disturbance related to vegetation diversity?

Although numerous case studies have documented the characteristics of swidden landscapes, it remains unclear whether spatial patterns exhibit any regularities across regions that differ in their ecological and cultural contexts. To address this gap, we extend previous work from southern Belize which found that the greatest levels of vegetation diversity tended to occur where there was some amount of anthropogenic disturbance [21]. This result is consistent with the intermediate disturbance hypothesis (IDH) [22], and aligns with studies which suggest a nonlinear relationship between disturbance and diversity [23, 24]. Here we examine whether this nonlinear relationship is evident in a sample of 18 swidden mosaics worldwide. Our analysis quantifies how variation in disturbance extent and configuration relates to vegetation diversity. Rather than seeking universal outcomes or rules pertaining to swidden disturbances, our goal is to identify broad regularities that link anthropogenic disturbance to spatial heterogeneity in swidden systems.

## 2 Methods

### 2.1 Sampling and Data Acquisition

We reviewed the ethnographic literature and identified 18 sites in the Central and South American Neotropics, equatorial Africa, and southeast Asia. These include sites in Belize (BLZ) [21, 25]; Brazil (BRA) [26]; Ecuador (ECU) [27, 28]; French Guiana (GUF) [29]; Nicaragua (NIC) [30]; Peru (PER) [31, 32]; Central African Republic (CAF) [33, 34]; Democratic Republic of Congo (COD) 1 [35, 36] and 2 [37, 36]; Madagascar (MDG) [10, 38]; Nigeria (NGA) [39]; Tanzania (TZA) [40]; India (IND) [41]; Indonesia (IDN) 1 [5, 42] and 2 [43]; Laos (LAO) [44]; Philippines (PHL) [45]; and Sri Lanka (LKA) [46, 47]) where swidden is known to be practiced. We sampled 4-band remotely sensed Planetscope imagery from Planet Labs [48] from a 50 km^2^ circular areas of interest around known swidden societies to encapsulate the surrounding swidden mosaics. We analyzed this imagery at down-sampled resolution of 9.375 meter pixels. In our reporting of these sites, we refer only to the nation state to protect the confidentiality of the residents of these locations.

### 2.2 Classification

**Landscape Classification.** We used supervised machine learning to classify each pixel as 1 of 13 classes (bare ground, recent clearing, early fallow, late fallow, mature forest, village/road, water, clouds, miscellaneous mask, palm oil fields, cattle pasture, urban, and wetland) using training data manually curated in QGIS [49].^1^ With this training data, we classified vegetation using a random forest model [50] and segment anything model [51]. The quality of classifications were assessed quantitatively using confusion matrices and accuracy metrics (Online Resource 1), and qualitatively by inspecting classifications for errors or artifacts. With the classification complete, we focus our analysis on the structure on swidden vegetation classes 1-5 (Figure 1).

**Figure 1:**
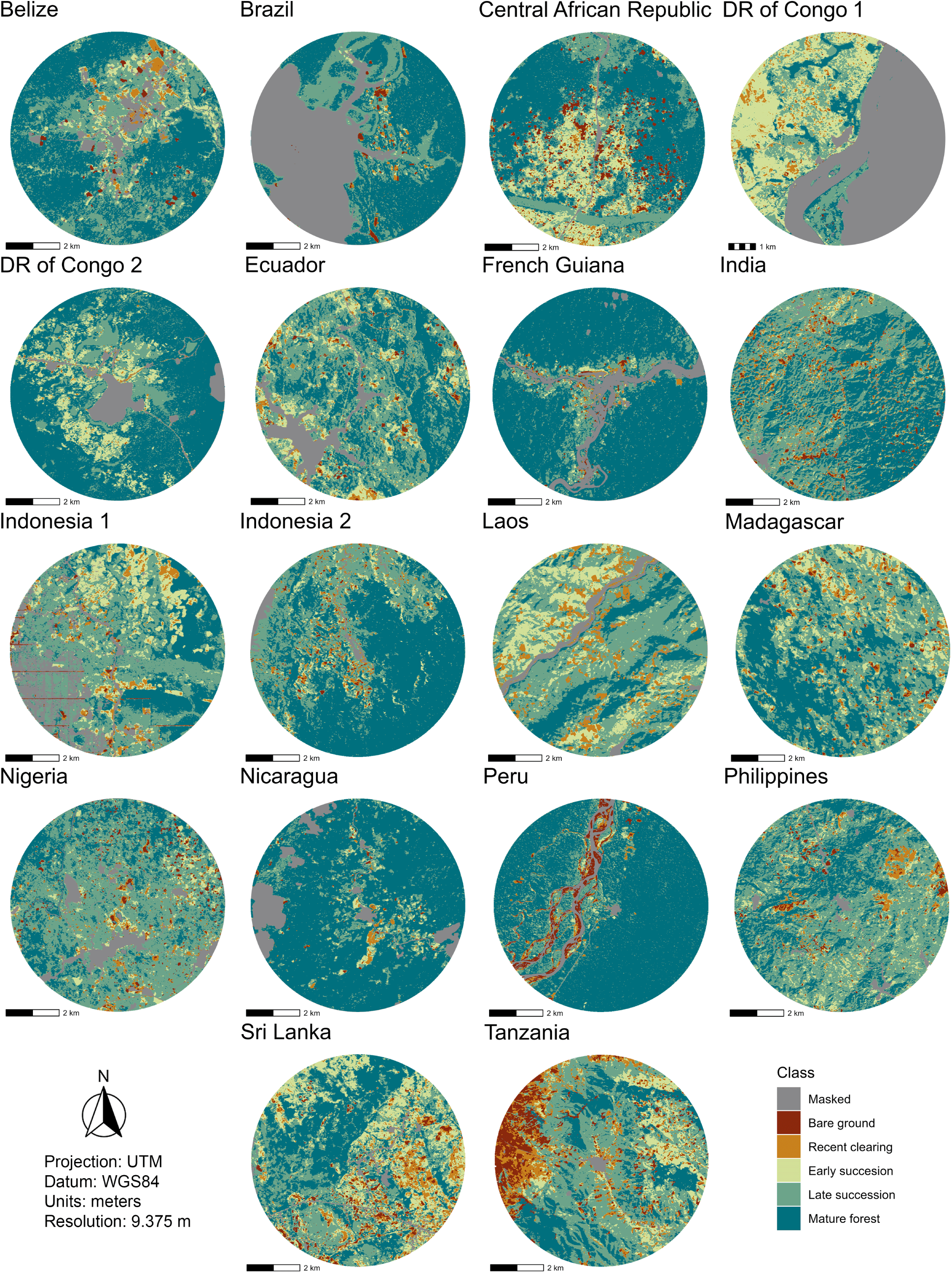
Classified m_3_a_0_ps of all 18 study sites.

**Spectral Species Classification.** We use spectral diversity as a proxy for vegetation diversity [52]. These signatures are detected through a process of radiometric filtering, dimensionality reduction (e.g., Principal Component Analysis), and spectral clustering (e.g., K-means) applied to multi-spectral imagery. We used the *biodivmapr* package [53] to classify each pixel in our raster as a one of 20 spectral species. Additional details and validation can be found be found in the Supplementary Information of [21].

### 2.3 Landscape Metrics and Covariates

To analyze the variability in the structural characteristics of swidden landscapes (Research Question 1), we used the *landscapemetrics* R package [54] to calculate a parsimonious set of landscape metrics to describe landscape structure. Based on recommendations in Cushman et al. [55], we selected a set of metrics that capture edge contrast, patch shape variability, contagion, juxtaposition, large patch dominance, and nearest neighbor distance/proximity. A full description of each metric and an ecological interpretation in the context of swidden is provided in the supplementary materials. To provide a minimal description of how landscape metrics vary across swidden societies, we apply a Principal Components Analysis (PCA), focusing on the first three axes of variation.

We examine local patterns of landscape structure by sampling the classified and spectral diversity maps using a hexagonal grid overlaid on the landscape using the *sf* package [56]. Our grid uses 2.5km wide cells, making each of them large enough to encompass an active swidden field and surrounding vegetation classes. We calculate disturbance *D*_i_ as composite measure of the degree of local patch density *P*_i_ and proportion of mature forest *F*_i_ that ranges from 0 (low disturbance) to 1 (high disturbance). *P*_i_ = 0 indicates that a grid cell is composed entirely of mature forest and *P*_i_ = 1 indicates extreme patch density with no mature forest present. As a measure of vegetation diversity *V*_i_, we calculate Shannon’s Index [57] for the spectral species within each cell and convert this value to effective species diversity (*e*^V^*^i^* ) which yields values that can be interpreted similarly to species richness [58].

### 2.4 Disturbance-Diversity Model

We model how anthropogenic disturbance relates to vegetation diversity in each cell *i* and site *u* using a multilevel Bayesian model. The full model specification and a discussion of priors can be found in supplementary materials. We model vegetation diversity *V*_iu_ as normally distributed with mean *µ*_iu_ and standard deviation *σ*. Because the spectral species classification procedure and our calculation of spectral species diversity described in section 2.3 produces strictly positive diversity values that cannot exceed 20, the mean *µ*_iu_ is obtained by transforming a latent predicted value *γ*_iu_ with a scaled logistic link function:

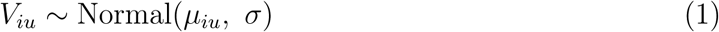

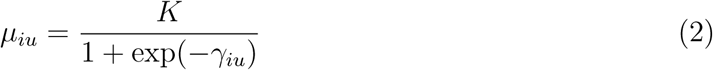

where *K* = 20 is the maximum possible number of spectral species. More generally, this link can be used for any upper bound *K*, and when *K* is not known, it can be estimated with a weakly informative prior. The effect of disturbance on vegetation diversity is represented by a quadratic function of disturbance:

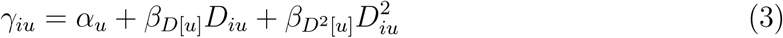

The parameters *β*_D[u]_ and *β*_D_2_[u]_ describe the shape of the disturbance-diversity curve for each site *u*. Allowing these slopes to vary by site partially pools information across sites, yielding both site-level curves and an overall (global) disturbance-diversity curve.

In the “Analytic Results” section of Supplementary Materials, we formalize a subset of the parameter space which would indicate empirical support for the IDH. These results identify the disturbance values *d*_1_ = 0.2 and *d*_2_ = 0.8 such that a unimodal curve that opens downward must peak between *d*_1_ and *d*_2_ to support the IDH. This formalism is generic and can be applied in future studies of disturbance effects on landscape mosaics.

All models were fit using the *brms* R package [59] and sampled using Markov Chain Monte Carlo with a no U-turn sampler implemented in Stan [60]. Using a step size of 0.99 and a tree depth of 12, all models converged with no divergent transitions on 4000 iterations and 1000 iteration warm-up period. *R*^^^ *<* 1.01 for all chains indicating that chains were properly mixed and bulk effective sample sizes show that the parameter space was adequately sampled.

## 3 Results

### 3.1 Structural Components across Swidden Mosaics

The *contagion* component characterizes the overall aggregation of vegetation classes across the landscape. Neotropical sites exhibit consistently high contagion values that reflect the presence of large, contiguous regions of mature forest. Similar patterns can be observed in selected sites in Indonesia and the Democratic Republic of the Congo. Patch density (Figure 2) serves as a compliment to contagion by providing a measure of the number of patches per unit area. Greater patch densities indicate greater levels of fragmentation and a lower degree of aggregation, a pattern that is strongest in Southeast Asian sites (Sri Lanka, India, Philippines, Indonesia 1), as well as some neotropical (Peru, Ecuador) and African (Central African Republic, Democratic Republic of Congo 2, and Nigeria) sites.

**Figure 2:**
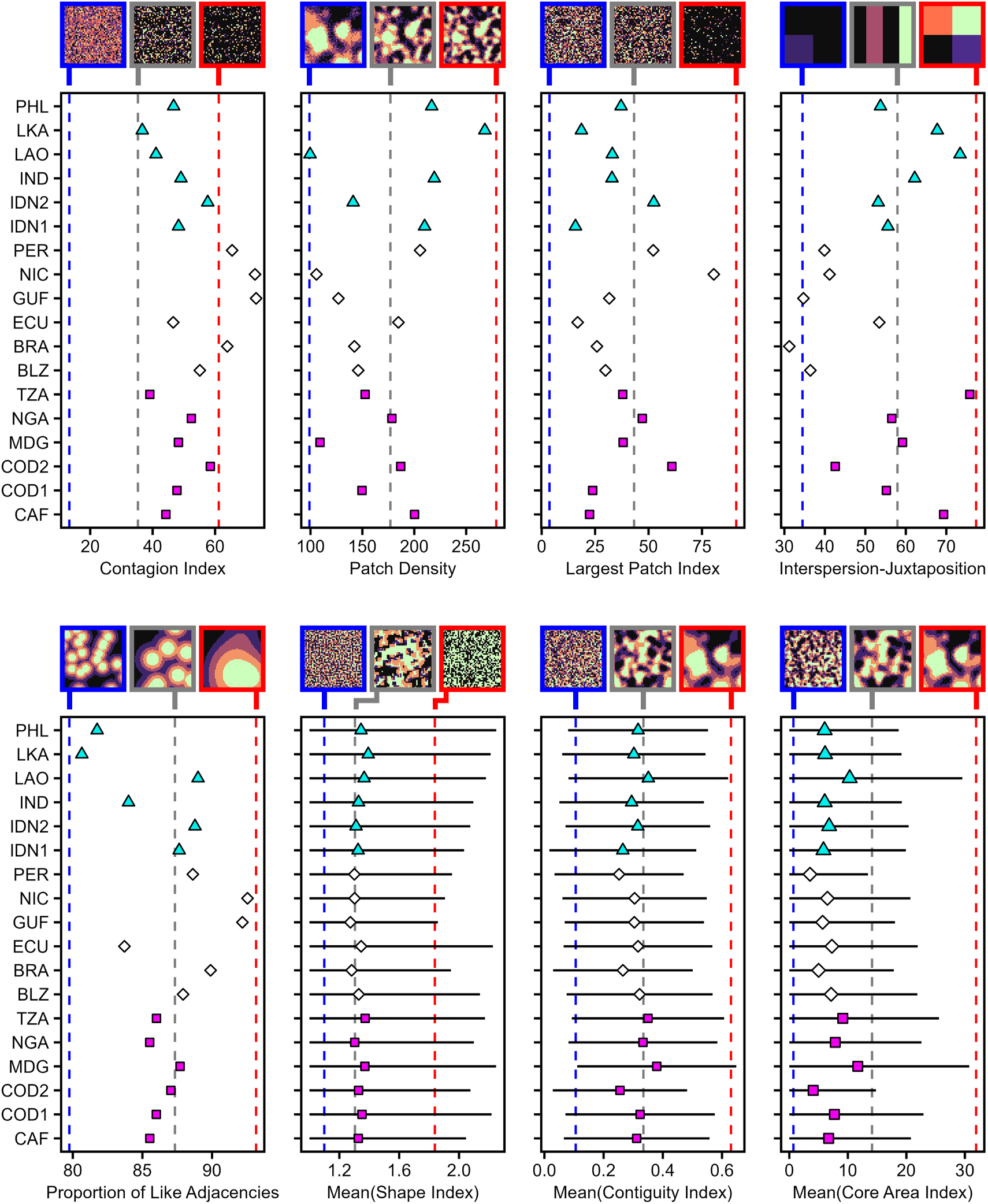
Landscape metrics describing the components of spatial mosaics across all 18 sites. Points show the computed landscape-level values (magenta squares = Africa; white diamonds = Neotropics; cyan triangles = Southeast Asia) and, where applicable, bars indicate one standard deviation above and below the mean. Dashed lines mark reference values derived from the example spatial patterns used heuristically to illustrate how metrics vary with different mosaic configurations.

The *large patch dominance* component captures the extent of a landscape that is concentrated within a small number of large vegetation patches. This is quantified by the largest patch index (Figure 2), which measures of the percentage of the landscape occupied be the single largest patch. Nicaragua stands out with 80% of the landscape dominated by a single mature forest patch, whereas other sites vary between 16% and 60%.

The *juxtaposition* component is measured using the interspersion and juxtaposition index (IJI, Figure 2), which quantifies how evenly patch adjacencies are distributed. This measures is analogous to a normalized entropy metric where greater values of IJI are associated with landscapes with more evenly distributed and intermixed classes. For example, the Nigeria site exhibits high IJI arising from a landscape composed of small forest fragments embedded within a diverse matrix of early successional, late successional, and mature forest patches. IJI tends to be lower in Neotropical landscapes due to the presence of larger, more aggregated patches.

The *edge contrast* component captures how frequently adjacent pixels belong to different vegetation classes. Rather than the magnitude of vegetation differences between adjacent patches, we focus on how often locally contrasting boundaries occur. We quantify this using the proportion of like adjacencies (PLADJ, Figure 2), which measures the fraction of all neighboring pixel pairs that are of the same class. Greater values of PLADJ indicate landscapes in which contrasting edges are uncommon, consistent with more aggregated and uniformly organized class configurations, whereas lower values suggest a greater prevalence of locally contrasting boundaries. Across all sites, PLADJ values range from 80% to 93%, indicating that swidden mosaics exhibit relatively few contrasting boundaries.

*Patch shape variability* characterizes variation in patch geometry and internal patch structure. Boundary complexity can be quantified using the mean shape index (Figure 2), which compares the patch perimeter to the perimeter of a square with equal area. Mean shape index ranges from 1.27 to 1.39, indicating that most patches are relatively compact and regular in shape. Greater values of the shape index indicate irregular shapes. Shape variability is greatest in Madgascar, Ecuador, and Democratic Republic of Congo 1; sites which all contain contiguous regions of late successional and mature forest. Mean contiguity reflects within-patch connectedness and mean core area index captures interior vegetation after accounting for edge pixels, so moderate contiguity suggests some spatial connectivity while low and highly variable mean core area implies many small patches embedded within broader vegetation matrices.

The *proximity* component captures the degree of isolation among patches of same vegetation class. Here, we use the mean nearest neighbor distance, where low values indicate relatively little isolation and a tendency of patches of the same class to occur close together in space. *Proximity* is complimented by the *nearest neighbor* component which captures the degree of patch dispersion. We measure this component using the standard deviation of nearest neighbor distances, which reflects variability in inter-patch spacing. Southeast Asian sites exhibit low variation in nearest neighbor distance, consistent with more regular spatial spacing, whereas Neotropical sites show substantially greater variation, indicating more spatially clustered patches.

#### 3.1.1 Axes of Variation

We conducted a principal components analysis of the landscape metrics described in Section 2.3focusing on the first three axes, which together explain 88.9% of the total variance (Figure 3.1.1). The first axis contrasts metrics associated with aggregation (contagion index, proportion of mature forest, largest patch index) with metrics describing configurational complexity and patch interspersion (IJI, mean shape index, and shape variability). Landscapes lying toward the aggregated end of this gradient are characterized by large contiguous forest patches surrounding fields with regular, compact shapes, frequently aligned along rivers and roadways. Examples of such mosaics are French Guiana and Nicaragua (Figure 1). The landscapes lying at the opposite end exhibit highly interspersed mosaics with irregular patch geometries, exemplified by Madagascar and Sri Lanka.

**Figure 3:**
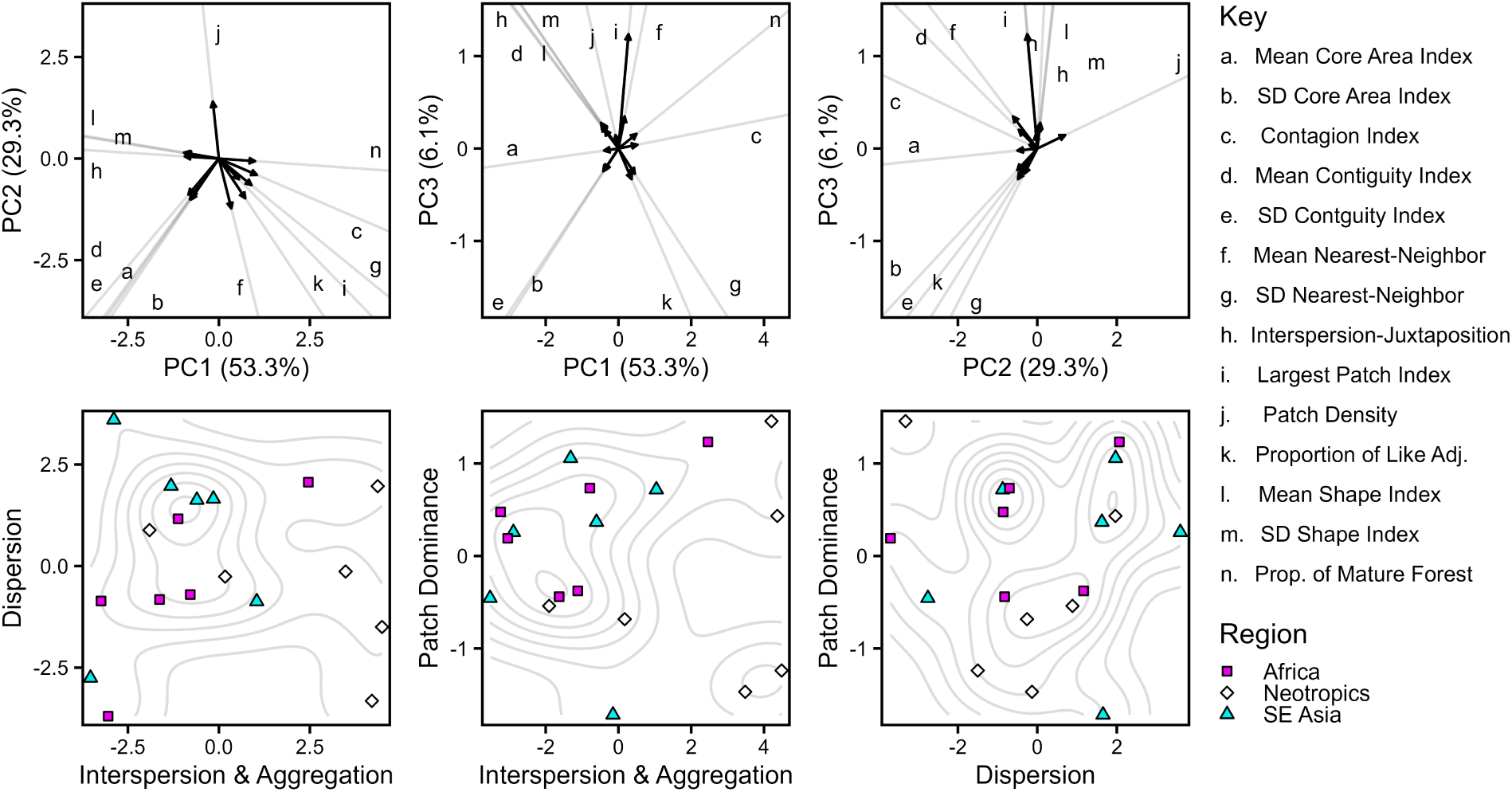
The first three PCs in a multivariate analysis of the landscape metrics. The top row illustrates the PC loadings, with variables labeled using lower case letters along corresponding grey reference lines. The bottom row illustrates sites projected into PC space. Contours in the bottom panels show the density interpolated using a Gaussian smoothing kernel.

The second axis is defined by a contrast between patch density, which loads positively, and both mean nearest-neighbor distance among same-class patches and the proportion of like adjacencies, which load negatively. This axis captures variation in the spatial dispersion versus proximity of patches of the same class, a gradient which reflects differences is how disturbance patches are spread across the landscape. Landscapes scoring toward the negative end of this axis are characterized by new clearings that are more widely spaced and spatially isolated (Figures 1 and 3). At the other end are landscapes which exhibit high patch density with short distances between neighboring patches, a configuration that arises when new clearings occur in close proximity with minimal buffers. Because swidden disturbances are relatively short-lived, such spatial proximity implies a partial temporal overlap in disturbance activity, which could be consistent with greater spatial synchronization. Sri Lanka illustrates this pattern, combining high levels of IJI on the first axis with locally concentrated disturbance activity on the second.

The third axis captures variation in spatial connectivity that is largely orthogonal to the dispersion gradient described by the second axis. While metrics related to internal patch structure and local connectivity (e.g., mean contiguity index and mean core area index) contribute to this dimension, the axis is defined primarily by the largest patch index. This indicates that an important component of variation across swidden mosaics is whether connectivity is achieved through the dominance of a single, large patch or through multiple, contiguous patches that are spatial intertwined. Landscapes like Nicaragua, Brazil, and Peru exemplify the former configuration, where connectivity is driven by dominant mature forest matrix, whereas other sites exhibit connectivity arising from several high-contiguity patches. In most cases these correspond to mature forest, although a subset of sites (e.g., Madagascar and Laos) also contained extensive and irregular regions of late successional forest.

### 3.2 Disturbance Curves

Our hierarchical model estimates distinct disturbance curves and diversity patterns for each landscape by partially pooling information from across sites, thereby improving both precision of our estimates and our understanding of the uncertainty of these relationships [61]. We can use these site level estimates to arrive at a global estimate which represents a general expectation about the shape of disturbance curves in swidden mosaics (Figure S3).

Figure S3 shows the average disturbance curve after marginalizing over the estimates from all 18 sites. At this aggregate level, the *β* parameters that dictate the shape of the disturbance curve indicate that, on average, the curve is unimodal and opens downward (*β*_D_ = 3.13 [SE = 0.56] and *β*_D2_ = −2.24 [SE = 1.03]). However, there is considerable variation in *β*_D2_ relative to *β*_D_ indicated by their respective *σ* parameters (*σ*_β_*_D_* = 0.79[SE = 0.76] vs. *σ*_β_*_D_*_2_ = 7.43[SE = 1.74]). The variation *β*_D2_ arises from sites with different curvature or which have fewer observations of extremely disturbed areas. These are identified in the next section.

Figure 4 shows the disturbance curves for each of the 18 sites. We provide 50 draws from the posterior predictive distribution to show a range of plausible relationships predicted by fitting data to equations 2-3. In the “Analytic Results” section of the Supplementary Materials, we define a region within which a unimodal peak of the disturbance curve would provide plausible evidence of the IDH. We observe such a peak in 12 of 18 sites. Indonesia 2 provides is a characteristic example of IDH curve, where the owest levels of disturbance are associated with moderate levels of vegetation diversity that increases at intermediate levels of disturbance, and declines to 0 as disturbance reaches the maximum.

**Figure 4:**
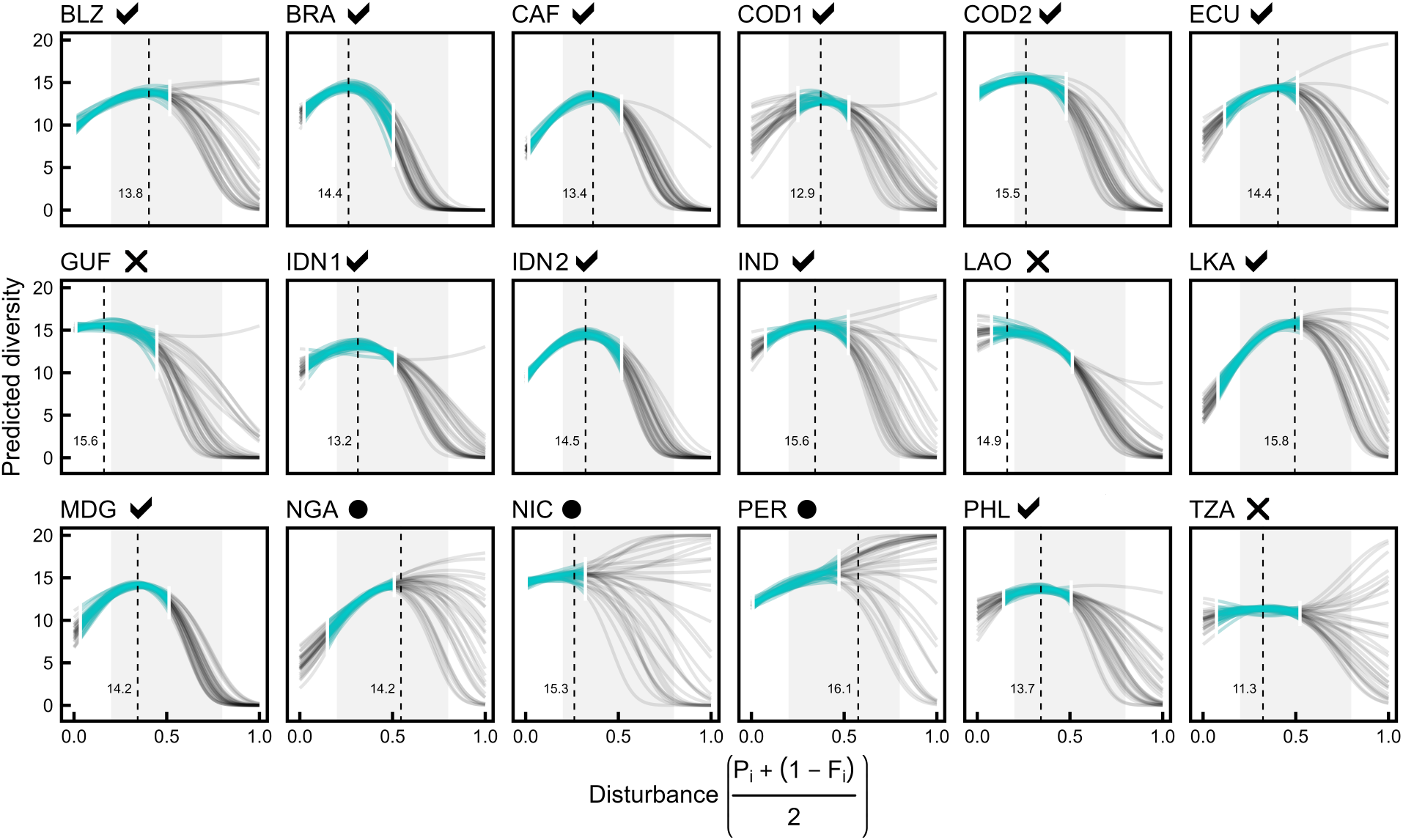
Predicted disturbance curves for all 18 swidden mosaics. Each line is a draw from the posterior distribution. Cyan lines shows the data range and grey lines show the predicted curve. Dashed lines indicate the location of the peak (*D*^*^) on the average curve and the shaded region is defined by *d*_1_ and *d*_2_. Sites with a peak located within this region support the IDH (i.e. *D*^*^∈ [*d*_1_, *d*_2_]) and are denoted by a black check mark. Sites that do not support the IDH or which are inconclusive are denoted by a × or •, respectively.

Of the six sites that do not support the IDH, two (French Guiana and Laos) exhibit peak diversity at low disturbance which then declines as disturbance increases. A third site (Tanzania) shows a relatively flat relationship at low and intermediate disturbance that becomes inconclusive at high disturbance. The final three sites are all inconclusive due to high uncertainty surrounding the *β*_D2_ estimate (Nigeria, Nicaragua, and Peru). Both the Nicaragua and Peru sites show very low levels of disturbance in general, leading to uncertainty around the shape of the full disturbance curve, even after partially pooling information from the other sites.

## 4 Discussion and Conclusion

### 4.1 Common Mosaic Patterns

This study applies methods from landscape ecology to analyze 18 swidden mosaics and identifies recurring structural configurations across diverse cultural and environmental settings. These mosaics can be described along three dominant configuration axes: the degree of aggregation versus interspersion of patches, the spatial dispersion versus co-occurrence of disturbance events, and the mode by which spatial connectivity is achieve. Taken together, these axes suggest that swidden mosaics occupy a constrained but flexible region of landscape configuration space, shaped by trade-offs among aggregation, dispersion, and connectivity rather than by any single sociocultural, economic, or environmental driver.

The first pattern is found in sites embedded within large tracts of mature forest where new fields and residences are systematically organized around rivers or roads. These landscapes exhibit aggregated patch structures and low levels of interclass mixing, forming “fishbone” configurations similar to those reported by de Oliveira and Metzger [4]. Such arrangements can arise in regions where expansion into mature forest regions is concentrated around infrastructural access or natural boundaries. Longitudinal analyses of forest cover in Amazonia suggest that these configurations can support forest regeneration, particularly when fallow lands facilitate soil recovery and successional processes [62]. At the same time, these configurations have also been interpreted as an early-stage of transformation in which infrastructure-driven expansion begins to dominate the landscape and may precede more extensive ecological change [63].

At the opposite end of this axis are landscapes with extreme levels of patch interspersion and juxtaposition that lack vegetation corridors to support movement and spatial connectivity [64]. Pardini et al., for example, link biodiversity to patch size distributions, arguing that landscapes dominated by small and relatively invariant patch sizes can impede dispersal processes, potentially leading to sudden regime shifts. Our analysis refines this interpretation by distinguishing between highly interspersed landscapes that differ in the spatial dispersion versus synchronization of disturbance patches. This distinction suggests alternative pathways to class intermixing, where forest recovery and resilience depend on whether interspersion arises from dispersed disturbances or from tightly co-occurring disturbance events.

A second pattern features moderate to high connectivity among mature forest and late successional vegetation, with swidden clearings more randomly dispersed throughout the landscape. Greater nearest-neighbor distances and a matrix of regenerating vegetation typify these sites, resembling the “independent settlement” pattern described by de Oliveira and Metzger [4]. Traditional ecological knowledge and cultural practices may contribute to such arrangements, particularly if rituals, norms, or social institutions encourage dispersed field placement strategies [42]. Theoretical modeling has also shown that landscapes which experience more randomized disturbances with limited aggregation exhibit a faster recovery [65] and are less likely to cause local population extinctions [66]. These insights suggest that we should be cautious of land-use policies that may intentionally or inadvertently encourage more aggregated spatial configurations [67, 68].

The third configuration includes 8 sites characterized by short interpatch distances and intraclass patches that co-occur in space. Such patterns can arise from myriad sociocultural, political, or market constraints such as competition for space by alternative land uses, labor scarcity, or policy environments that inhibit forest clearance [67]. In Sri Lanka, for instance, forest clearing restrictions that promote compact field arrangements; in Indonesia 1 and Belize, expanding oil palm and cattle enterprises may constrain available land. The Sri Lanka case in particular demonstrates the complex interaction of smallholder farm activity, land use policy, and conservation strategies, often centered around human-wildlife conflicts and development agendas [69]. Anuradha et al [69], for example, argue that the transition from swidden cultivation to fixed, continuous agriculture has been associated with a substantial increase in conflict that is not observed in regions with more traditional agricultural practices. Such conflicts are yet another indicator that managing forest landscapes requires a strategy that integrates people, wildlife, and ecosystems as part of a complex, socio-ecological system [70]. We believe swidden agriculture can serve a model system for investigating and understanding the implications of such strategies [6, 15].

### 4.2 Localized Disturbance Patterns and Vegetation Diversity

The measure of vegetation diversity used in this study provides a scale-appropriate indicator for examining vegetation diversity across large spatial extents that would be impractical to assess using detailed field inventories. The spectral diversity approach applied here has been interpreted as an integrated signature of functional and phylogenetic plant diversity, based on established links between spectral reflectance and variation in plant morphology [71, 52]. From this perspective, spectral diversity could reflect the degree to which functionally distinct plant-assemblages co-occur within local landscape units.

If swidden cultivation uniformly degraded forest landscapes, we would expect vegetation diversity to peak at low levels of disturbance and to decline monotonically as disturbance intensifies, though this pattern is rarely observed in our analysis. The closest example occurs in Laos, where high connectivity among successional vegetation coincides with low proportions of mature forest and low patch density, producing large, aggregated disturbance areas. Aggregated successional classes may limit dispersal by increasing average distances over which propagules must disperse to regrow into recently abandoned fields within a matrix with diminished seed-bank potential [72, 65]. We also do not observe a unimodal disturbance-diversity relationship in French Guiana, where disturbances are strongly concentrated along river corridors. Despite the high diversity associated with the surrounding mature forest matrix, disturbed patches in this site tend to have limited adjacency to mature forest. This suggests that pioneering clearings could increase functional heterogeneity over time by integrating disturbances into a more diverse forest matrix [73].

Across most of the landscapes examined, we observe unimodal disturbance-diversity relationships consistent with the IDH suggesting that nonlinear responses of vegetation diversity to anthropogenic disturbance are a common feature of swidden mosaics. Rather than indicating a universal mechanism, a more cautious interpretation is that unimodal disturbance-diversity relationships emerge regularly within the constrained configuration space identified by our multivariate analyses. This persistence suggests that the disturbance-diversity relationship is not solely determined by overall landscape configuration or aggregate levels of disturbance, but instead reflects interactions between localized disturbance and the surrounding spatial context.

Taken together, these findings emphasize the importance of understanding how anthropogenic swidden mosaics contribute to the vegetation diversity of tropical landscapes. Such contributions cannot be inferred from aggregated rates of land clearing alone. Instead, they likely depend on how disturbances are spatially arranged and the sociocultural and economic mechanics that shape field placement and size. Previous work in Belize illustrates how local customs and traditional ecological knowledge can shape disturbance placement and produce heterogeneous landscape structure that have the potential to benefit vegetation diversity and productivity [74, 21, 75]. Integrating such fine-grained social and ecological processes with landscape-scale modeling offers a promising avenue for evaluating and refining the macro-scale predictions presented in this study.

### 4.3 The Implications of Swidden as a Disturbance Regime

While comparing swidden to wet-rice terraces in Java, the ecological anthropologist Clifford Geertz observed that agricultural systems differ not only in productivity but how they restructure ecosystems:

“Any form of agriculture represents an effort to alter a given ecosystem in such a way as to increase the flow of energy to man: but a wet-rice terrace accomplishes this through a bold reworking of the natural landscape; a swidden through an uncanny imitation of it” [76, p. 16].

Subsequent research in ecological anthropology and agroecology has reinforced this view, noting that swidden and other agroecosystems can mimic key structural features of tropical forests [13, 77]. Much of this work has focused on intercropping, stratification, and successional cycling that parallels forest ground-cover, understory, and canopy structure. We add to this line of argument by examining how the spatial configuration of anthropogenic disturbance shapes landscape-level patterns.

From this spatial perspective, our results suggest that swidden disturbances are not an external, uniform disruptor of forest dynamics. Across most sites, vegetation diversity peaks at intermediate levels of disturbance, consistent with disturbance-mediated coexistence mechanisms emphasized in the IDH [22]. In this view, disturbance appears to reduce the dominance of competitively superior species while preserving sufficient connectivity and vegetation extent to support dispersal and recovery processes, creating a set of conditions under which disturbance could theoretically enhance diversity [78]. Rather than treating swidden has inherently degradative, these findings support a reframing of swidden as a disturbance regime whose ecological effects can mimic those of natural disturbance processes under certain scales and configurations.

These insights have direct implications for land-use governance, conservation policy, and our general understanding of the role of anthropogenic disturbance in forest ecosystems. Management strategies that confine swidden to narrow corridors, roadside settlements, or fixed enclosures may inadvertently simplify a landscape, reducing spatial heterogeneity and undermining some of the ecological processes that sustain diversity at the landscape-scale. Such constraints can also limit community agency and disrupt long-standing relationships between cultural practices, land use, and forest dynamics. From a landscape perspective, this suggests that policies aimed at reducing deforestation or promoting conservation must consider not only the extent of anthropogenic disturbance, but also its spatial composition, configuration, and the ways that culture mediates such patterns.

Tropical forests have been shaped by agricultural practices for millennia [79, 80]. Human modification of forests began tens of thousands of years ago [81], and recognizing this deep history is essential for contemporary conservation and restoration efforts [82]. The consistent emergence of unimodal disturbance-diversity relationships across swidden mosaics examined here suggests that anthropogenic disturbances may be integral components of tropical forest ecosystems. This perspective aligns with a growing recognition that human activities are not strictly external to ecological systems, but are embedded within them [83]. Advancing this view requires analytical approaches that recognize socially mediated disturbance and landscape heterogeneity as elements of a complex adaptive system [84]. Swidden systems, in this sense, serve as a valuable model system for understanding how long-term human-environment interactions shape ecological resilience in a rapidly changing world.

## Acknowledgements

We thank Bill Peterman for his comments and advice related to this manuscript. We also thank Simon Nabors for his early contributions to the machine learning pipeline used to classify these landscapes. We extend our appreciation the staff at Planet Labs who answered key questions related to the spatial imagery used in this study.

## Footnotes

1 For community confidentiality, certain features in Figure 1 were redacted for publication. All analyses were conducted on unredacted datasets.

