## Supplementary Materials for "Anthropogenic disturbance and spatial heterogeneity shape vegetation diversity in tropical swidden mosaics globally"

February 27, 2026

### S1 Landscape Metrics and Their Interpretations in Swidden

We use landscape metrics [1] to quantitatively describe the patch geometry of swidden mosaics. Metrics can be calculated at the landscape, class, or patch level, where landscape refers to the entire raster object, class refers to distinct categories of vegetation within the landscape, and patch refers to a single instance of a vegetation class. Our approach follows the recommendations provided by [2] about the set of landscape characteristics needed to comprehensively describe a landscape. These are *contagion, large patch dominance, interspersion and juxtaposition, edge contrast, patch shape variability, proximity,* and *nearest neighbor distance.* Unless otherwise stated, all of our metrics are calculated at the landscape scale.

In this section, we define the metrics that we use to characterize these landscape characteristics [1]. We state the name of the metrics along with the code function (e.g., `lsm_l_contag`). All of the equations along with descriptions of the metrics and how to use them R are located on the `landscapemetrics` github website. Here we provide a brief description along with an ecological interpretation related to swidden.

#### S1.1 Edge contrast

**Proportion of Like Adjacencies** (`lsm_l_pladj`) – A like adjacency means that two cells of class \$i\$ are adjacent to one another, and is calculated as

$$\text{PLADJ} = \left(\frac{g_{ii}}{\sum\limits_{k=1}^{m} g_{ik}}\right) * 100 \tag{1}$$

where $g_{ii}$ is a count of adjacent pixels within a class and $g_{ik}$ is a count of the adjacent pixels between classes. Landscapes with a high aggregation or clustering and large contiguous patches are likely to have greater proportion of like adjacency (PLADJ). A chess board pattern, for example, would have PLADJ = 0 because each cell is only adjacent to unlike cells that are part of a different class (in the rook's case). PLADJ shares some information with measures like contagion which capture the degree of clustering in the landscape. Interestingly, all of our sites from Africa show very little variation in PLADJ, with all sites having PLADJ around 0.86.

#### S1.2 Patch Shape Variability

**Average Shape and Shape Variation** (`lsm_l_shape_mn`, `lsm_l_shape_sd`) – The shape index measures how much a patch deviates from a perfect square. The index is calculated as

$$\text{SHAPE} = \frac{0.25 * p_{ij}}{\sqrt{a}_{ij}} \tag{2}$$

where $p_{ij}$ is a patch and $a_{ij}$ is the area of that patch in $m^2$. When shape = 1, a patches are perfectly square and in our swidden mosaics, average shape ranges between 1.275 and 1.375, suggesting that most patches are square-like. This semi-regularity makes sense given that many patches arise from the creation of agriculture fields. However, variation in shape index is considerable across all sites. High values of shape index indicate more complex shapes, which arise where there are more complicated edges between adjacent patches, perhaps as vegetation successes after disturbance. Such complicated edge pattern may provide valuable microenvironments and new niche fulfillment opportunities.

**Contiguity** (`lsm_l_contig_mn`, `lsm_l_contig_sd`) – The contiguity index (CONTIG) is a different kind of shape index which captures the degree to which pixels in a patch are highly connected. The index is calculated as

$$\text{CONTIG} = \frac{\left[\frac{\sum\limits_{r=1}^{z} c_{ijr}}{a_{ij}}\right] - 1}{v - 1} \tag{3}$$

where $c_{ijr}$ is the contiguity of each pixel within a patch, which is calculated in reference to the area $a_{ij}$ of two patches $ij$. The size of the filter window $v$ is also used to constrain the contiguity values, though we only use default settings. CONTIG can serve as a measure of the shape of patches, and in particular, the overall connectivity of patches in a landscape. This measure was first introduced by [3] who believed that the contiguity index would aid in management goals geared toward improving the connectivity of similar habitats. Thus, swidden landscapes which have high contiguity can interpreted as having high connectivity. As structural features, these connected patches are important for dispersal and movement, depending on the vegetation class that exhibits the contiguity.

### S1.3 Contagion/Diversity

**Contagion** (`lsm_l_contagion`) – The contagion index captures the probability that two random chosen adjacent pixels belong to the same class and is calculated using a weighted entropy similar to Shannon's Index

$$\text{CONTAG} = 1 + \frac{\sum\limits_{q=1}^{n_a} p_q ln(p_q)}{2ln(t)} \tag{4}$$

Here $p_q$ is a cellwise adjacency table across all vegetation classes and $t$ is the total number of classes in the landscape. The numerator of this equation is the weighted entropy, which is rescaled based on the weighted probability of each landscape class. Contagion is often interpreted as a measure of the "clumpiness" or the clustering of pixels within a landscape, but more generally it provides a measure of spatial information. As pointed out by [4], this measure is influenced by how evenly distributed the classes are. If there is one dominant class, this can drive up the measure of CONTAG just as a high abundance of a single dominant species can drive up the value of a Shannon Index when it is used to calculate species diversity [5]. For example, contagion is high in our Nicaragua site because the village in situated within a large expanse of mature forest. For this reason, it is useful to compare contagion to the largest patch index (LPI) or total class area. Indeed, our own analysis shows that contagion is closely related to the proportion of the landscape covered in mature forest. If mature forest patches form large clusters, this can drive greater values of the contagion index. A landscape with a high proportion of mature forest and high contagion but without a high LPI usually arises when the area of interest (AOI) is divided by a river or road, leading to a smaller number of large forest clusters. However, we expect that low contagion, particularly when the proportion of forest is low, could cause ecological issues because the swidden mosaic may lack a core area that serves as a seedbank for dispersal.

### S1.4 Juxtaposition

**Core Area Index** (`lsm_l_cai_mn`, `lsm_l_cai_sd`) – Any single patch of vegetation has what are known as core and periphery pixels. The peripheries are those pixels in a raster that lie at the edge between two classes. The core area, then, is composed of the pixels which are only surrounded by pixels of the same class within a patch (see [1]); or, alternatively, those pixels which do not lie adjacent to another class. Holding area constant, the shapes which have the largest core area would be squares and circles. For example, given a square and a rectangle with the same area, a square will have a greater core area than the rectangle, because fewer pixels would lie along the perimeter.

The *core area index* (CAI) is a measure of the percentage of core area pixels relative to the area of a patch

$$\text{CAI} = (\frac{a_{ij}^{core}}{a_{ij}}) * 100) \tag{5}$$

Here $a_{ij}^{core}$ is the area of pixels that are part of the core. For a complex mosaic, a higher mean core area suggests the presence of many large patches with more regular shapes. Low CAI would index high fragmentation and/or the presence of long, winding patches, which would be indicated by high contiguity. These structures are ecologically significant because vegetation patches with a large core area, especially mature forests, are expected to have a large influence on seed dispersal because the vegetation within the core serves as a seed bank that is less affected by biotic and abiotic factors that may be stronger along the edge. Long and winding patches (low core area) can be both beneficial and problematic. While these corridors may offer considerable connectivity, they may also be vulnerable to the spread of fires, pests, or invasive species (see [6] for a discussion of labyrinthine Turing patterns in landscape structure). If the coefficient of variation is high, this indicates the presence of many patch sizes and structures, both regular and irregular.

**Interspersion and Juxtaposition Index** (`lsm_l_iji`) – [1] refer to this as a "salt and pepper" metric that describes the degree to which patches of different classes are intermixed. This index is calculated as

$$\text{IJI} = \frac{-\sum_{i=1}^{m}\sum_{k=i+1}^{m}\left[\left(\frac{e_{ik}}{E}\right)ln\left(\frac{e_{ik}}{E}\right)\right]}{ln(0.5[m(m-1)])} * 100 \tag{6}$$

where $e_{ik}$ are the upper and lower triangles of the adjacency matrix of all classes. $E$ is the total edge length and $m$ is the total number if unique classes. IJI contains an expression that is similar to Shannon's Entropy, However, the value of entropy is normalized by the maximum possible entropy given the number of classes in the landscape. IJI is necessarily low in sites that have large patches of a single class (e.g., mature forest). IJI can also be low if classes are of relatively equal size and patches of vegetation class tends to occur near to one other This happens in highly ordered ordered landscapes. Classes which intermix but have a large patch size relative to the scale of the landscape can still lead to a relatively high IJI index.

### S1.5 Large Patch Dominance

**Largest Patch Index** (`lsm_l_lpi`) – The largest patch index (LPI) goes hand in hand with other measures of landscape structure and configuration. It is simply the percent of a landscape covered by a single patch, regardless of the class of that patch

$$\text{LPI} = \frac{\max(a_{ij})}{A} * 100 \tag{7}$$

In our analysis, sites with high LPI tend to be those with a large contiguous region of mature forest (e.g., Nicaragua). Nigeria stands out as a site with an LPI of almost 50% with a low proportion of the landscape covered in mature forest. In the Nigeria case, the largest patch is a secondary forest that largely surrounds the village center and roadways. Indonesia 1 – a site with a large amount of fragmentation from swidden and palm oil production – also stands out with a very low LPI.

**Proportion of mature Forest** (`lsm_p_area`) – The proportion of mature forest (PF_prop) is the sum of the area of all patches in the mature forest class (see also Variable Calculation). It is a highly important feature of landscape structure, and the proportion of mature forest is often used to estimated how disturbed a forest landscape is. In some sense, regions of mature forest represent the reservoir of seeds that can disperse and regrow into disturbed regions. Along with LPI and patch density (PD), PF_prop can tell us whether a landscape has many mature forest fragments.

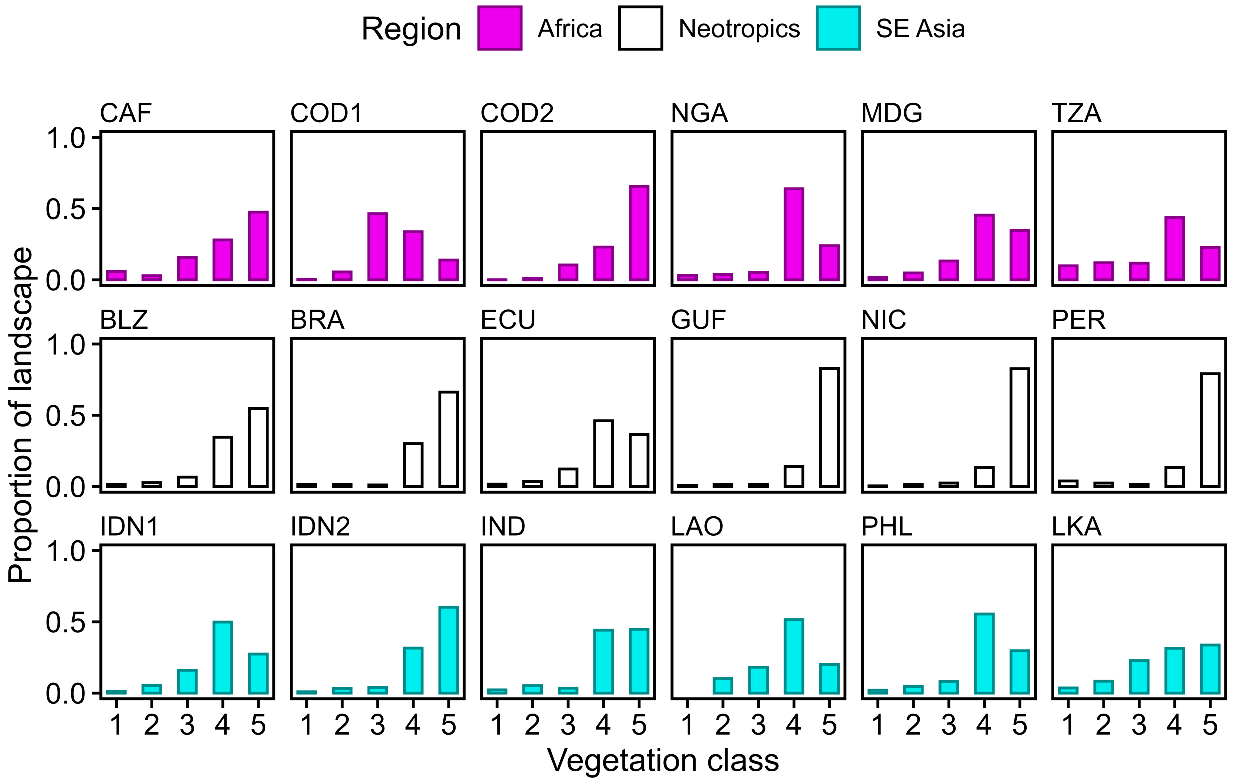


Figure S1: The proportion of each landscape covered by each vegetation class (1 = bare ground, 2 = recent clearing, 3 = early succession, 4 = late succession, 5 = mature forest).

Table S1: Summary of landscape metrics across all 18 landscapes.

| Metric | Units | Mean | Range |
|---|---|---|---|
| Mean Core Area Index | Percent | 6.86 | 3.54 - 11.70 |
| SD Core Area Index | Percent | 14.15 | 9.90 - 19.25 |
| Contagion | Percent | 52.59 | 36.67 - 73.14 |
| Mean Contiguity Index | None | 0.31 | 0.25 - 0.38 |
| SD Contiguity Index | None | 0.25 | 0.22 - 0.27 |
| Mean Nearest Neighbor Distance | Meters | 38.50 | 31.70 - 47.94 |
| SD Nearest Neighbor Distance | Meters | 44.44 | 27.28 - 67.42 |
| Interspersion-Juxtaposition | Percent | 53.41 | 31.22 - 75.79 |
| Largest Patch Index | Percept | 36.51 | 15.81 - 80.54 |
| Patch Density | N / 100ha | 169.07 | 99.60 - 268.04 |
| Proportion of Mature Forest | Proportion | 0.46 | 0.14 - 0.82 |
| Proportion of Like Adjcencies | Percent | 86.92 | 80.65 - 92.55 |
| Mean Shape Index | None | 1.33 | 1.27 - 1.39 |
| SD Shape Index | None | 0.77 | 0.58 - 0.90 |

#### S1.6 Nearest Neighbor Distance/Proximity

**Euclidean Nearest Neighbor Distance** (`lsm_l_enn_mn`, `lsm_l_enn_sd`) – Nearest neighbor distances (ENN) are calculated by computing the Euclidean distance between the centroids of patches in the same classes

$$\mathrm{ENN} = h_{ij} \tag{8}$$

where $h_{ij}$ is a Euclidean distance between patches $i$ and $j$. At the landscape level, these distances are averaged. Lower values of this metric indicate that patches tend to be close to one another in space. This serves as an indication of the degree of spatial synchronization. For example, in Sri Lanka, the average ENN is very low, suggesting that patches in a particular class tend to co-occur close to one another. Ecologically speaking, these aggregated patches can functionally serve a one large disturbance rather than several smaller disturbances, and this can limit the capacity of swidden to enhance the structure of a tropical forest landscape. When ENN is greater, patches are more spread out cross the landscape, suggesting that disturbances are more distributed more randomly in space.

**Patch Density** (`lsm_l_pd`) – Patch density (PD) is one of the simplest but most important measures of landscape structure. Calculated as the number of patches divided by the area of the landscape

$$\mathrm{PD} = \frac{N}{A} * 10000 * 100 \tag{9}$$

PD is greatest when each pixel in a raster is considered a separate patch. Although PD does not take into account the configuration of patches, it captures a very important component of disturbance. Landscapes with very high patch density are highly fragmented while those with very low patch density will likely also have high LPI. Sites with low PD are those which have large contiguous regions of mature forest, or those which have larger patches on average.

### S2 Model Description an Notation

In this section, we describe the variables used in the hierarchical model along with the model form, notation, and priors.

#### S2.1 Local Variable Calculations

To measure the local characteristics of vegetation structure, we impose a hexagonal grid over both the classification and spectral species maps. Within each sampling unit $i$ of this grid, we calculate focal landscape metrics [1] and summarize spectral species [7]. By changing the diameter of the sampling units, it is possible to study landscape structure at multiple scales within each swidden society.

We calculate Shannon Information (H) of the spectral species in each sampling unit as a measure of local vegetation diversity (i.e. beta diversity)

$$H_i = -\sum_{k \in \mathcal{K}} p(s_{ik}) \log_2 p(s_{ik}) \tag{10}$$

where $s_{ik}$ is a distinct spectral species $k$ within the set of possible species $\mathcal{K}$ in each sampling unit $i$ imposed on the landscape. We then convert this value to the effective number of species

$$V_i = e^{H_i} \tag{11}$$

where $V_i$ is a vector of continuous values between 0 and K (maximum spectral species clusters) that can be interpreted similarly to species richness [5]. This is the outcome variable of interest in our Bayesian models.

Within a sampling unit $i$, patches are labelled $c$ with the respective class where $c = 1, 2, ...13$ possible classes. We calculate the proportion of each sampling unit $i$ that is covered in mature forest ($F_i$) by summing the area of all patches labelled as mature forest and dividing by the total sampling unit area

$$F_i = \sum * j * \in \mathcal{J}_{\rangle \bigtriangledown} a_{ij5} / A_i \tag{12}$$

where $a_{ij}$ is the area of each patch $j$ of class $c = 5$ in sampling unit $i$, and $A_i$ is the total area of the sampling unit in $m^2$. Although $A_i$ will be the same for most sampling units, those that contain masked regions will have 'NA' values that reduce the value of $A_i$. Thus, $F_i$ is a proportional measure that is comparable across sampling grids $i$ with distinct areas because $0 \leq F_i \leq 1$ for all $i$. Moreover, we can compute $1 - F_i$ as the proportion of each sample unit that is not covered in mature forest, which serves as a more intuitive proxy for disturbance. For example, when $1 - F_i = 0$ the sample unit is entirely covered in mature forest, a state which we would intuitively described as undisturbed. As $1 - F_i$ increases, the sample unit experiences more disturbance.

While $1 - F_i$ gives a proxy of disturbance based on area, other measures capture the structure of disturbance, whether by edges or by fragmentation. We focus on patch density $P_i$ as a measure of disturbance intensity associated with fragmentation. We count the number of all patches $j$ within a cell and divide by the area

$$N_i = \sum_j 1 \tag{13}$$

$$P_i = \frac{N_i}{A_i} * 10000 * 100 \tag{14}$$

The patch density is expressed as the number of patches per 100 hectares. As the intensity of disturbance increases, patch density increases. We normalize $P_i$ between 0 and 1.

### S2.2 Single-Site Model Form

We model the vector $V_i$ using the Gaussian model family with mean $\mu_i$ and standard deviation $\sigma$

$$V_i \sim \text{Normal}(\mu_i, \sigma). \tag{15}$$

Given that $0 < V_i < K$ for all landscapes, we model $\mu_i$ using a scaled logistic function

$$\mu_i = \frac{K}{1 + \exp(-\gamma_i)} \tag{16}$$

where $\gamma_i$ dictates the geometry of the disturbance curve. The numerator $K$ is the asymptotic limit of the disturbance curve – the maximum value of vegetation diversity – which is determined by the clustering algorithm used in the spectral species classification (e.g. $K = 20$; see documentation for [7]).

Each measure comes from a grid cell in the overlaid hexagonal grid. We include sampling unit level variables using the following submodel

$$\gamma_i = \alpha + \beta_D D_i + \beta_{D^2} D_i^2 \tag{17}$$

where $\alpha$ estimates the average vegetation diversity (the y-intercept). We model the association of patch density and vegetation diversity using a 2nd-order polynomial by squaring $D_i$ and estimating parameters $\beta_D$ and $\beta_{D^2}$. $D_i$ represents a single composite measure of disturbance made of up $P_i$ and $F_i$

$$D_i = \frac{P_i + (1 - F_i)}{2} \tag{18}$$

We note that this composite measure is an average of the two normalized measures. This is an alternative to the typical $z$-score approach that provides the same results. We choose this alternative because it maintains the same 0 to 1 range which is easily interpreted as 0 = no disturbance and 1 = extreme disturbance. By taking $1 - F_i$, we can interpret the composite value 0 as a sample unit which is entirely composed of a single mature forest patch – a forest condition that is conventionally thought of as "undisturbed". As $D_i$ increases, the sample unit becomes more fragmented (higher patch density) and less composed of mature forest.

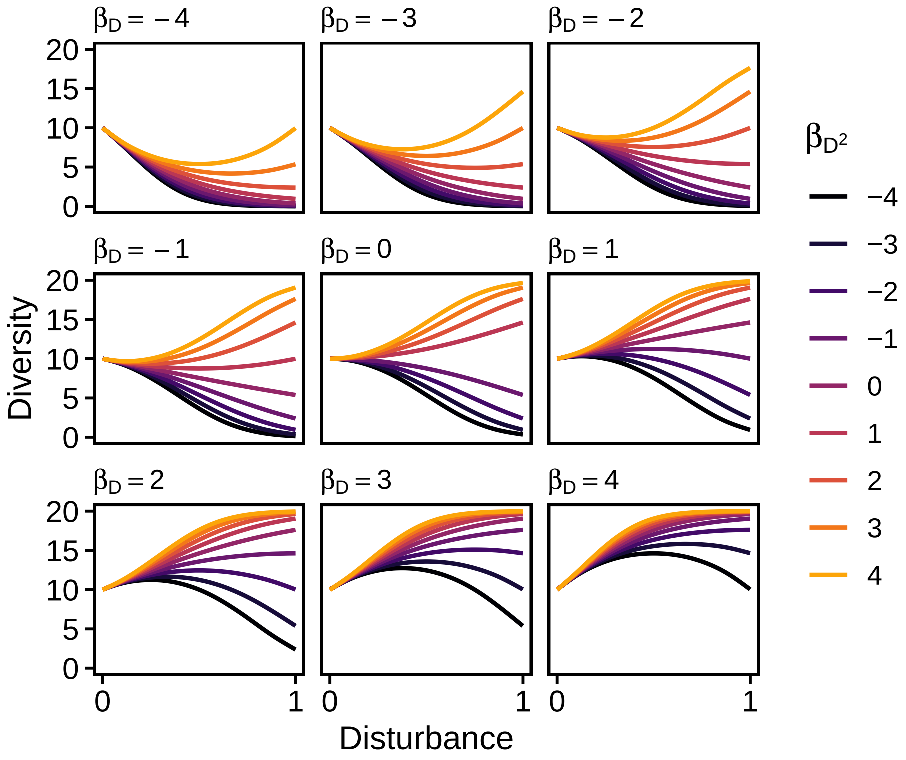


Figure S2: Examples of disturbance-diversity response curves generated by the model using different combinations of parameter values for $\beta_D$ and $\beta_{D^2}$.

#### S2.2.1 Full Single-Site Model with Priors

The full version of the model contains our composite measure along with all priors for model parameters.

$$V_i \sim \text{Normal}(\mu_i, \sigma) \tag{19}$$
$$\mu_i = \frac{K}{1 + \exp(-\gamma_i)} \tag{20}$$
$$\gamma_i = \alpha + \beta_D D_i + \beta_{D^2} D_i^2 \tag{21}$$
$$\alpha \sim \text{Normal}(0, 2) \tag{22}$$
$$\beta_D, \beta_{D^2} \sim \text{Normal}(0, 1) \tag{23}$$
$$\sigma \sim \text{Exponential}(1) \tag{24}$$

The priors are set as weakly regularizing. For example, the prior for all $\beta$ parameters is Normal$(0, 1)$ which is agnostic about the direction of these parameters – which allows many possible curves – but is skeptical about very strong associations (i.e., greater than 2 standard deviations above the mean). Given previous work [8], we could justify an informative prior in which $\beta_{D2}$ is negative, on average. However, for a single site model, we remain agnostic about the direction of this effect.

### S2.3 Hierarchical Model

We fit a multilevel model across all sites treating each site $u$ as a cluster of varying effects. This makes it possible to fit a unique disturbance curve for each site based on partial pooling of variation across all $i$ within all sites. We can then estimate a global disturbance curve by marginalizing over all sites.

**Data model**

$$V_{iu} \sim \text{Normal}(\mu_{iu}, \sigma) \tag{25}$$
$$\mu_{iu} = \frac{K}{(1 + \exp(-\gamma_{iu}))} \tag{26}$$
$$\gamma_{iu} = \alpha_u + \beta_{D[u]} D_{iu} + \beta_{D^2[u]} D_{iu}^2 \tag{27}$$

**Group $u$-level nonlinear parameters**

$$\begin{bmatrix} \alpha_u \\ \beta_{D[u]} \\ \beta_{D^2[u]} \end{bmatrix} \sim \left( \begin{bmatrix} \bar{\alpha} \\ \bar{\beta_D} \\ \bar{\beta_{D^2}} \end{bmatrix}, \quad \mathbf{\Sigma} \right) \tag{28}$$
$$\mathbf{\Sigma} = \mathbf{DRD} \tag{29}$$
$$\mathbf{D} = \text{diag}(\sigma_\alpha, \sigma_{\beta_D}, \sigma_{\beta_{D^2}}) \tag{30}$$

**Hyperpriors**

$$\bar{\alpha} \sim \text{Normal}(0, 2) \tag{31}$$
$$\bar{\beta_D} \sim \text{Normal}(0, 1) \tag{32}$$
$$\bar{\beta_{D^2}} \sim \text{Normal}(-1, 1) \tag{33}$$

**Standard deviations**

$$\sigma_\alpha, \sigma_{\beta_D}, \sigma_{\beta_{D^2}}, \sigma \sim \text{Exponential}(1) \tag{34}$$

**Correlation structure**

$$\mathbf{R} \sim \text{LKJCorr}(\eta), \qquad \eta = 4 \tag{35}$$

The priors represent weakly informative and regularizing priors. For example, we set all $\sigma$ priors to Exponential(1), which is skeptical of large deviations while still accommodating large deviations. We also set the prior for $\bar{\beta}_{D2}$ to Normal$(-1, 1)$ based on our prior simulations and our expectations from previously published work [8]. Despite this being an expectation of a unimodel disturbance curve that opens downward, with multiple sites and repeated measure within sites, it leaves plenty of room for functional forms that do not support the IDH.

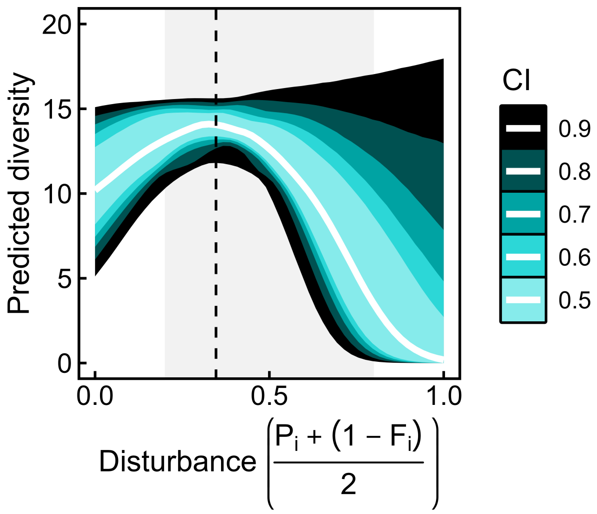


Figure S3: The average disturbance curve after marginalizing over parameter estimates from all 18 sites. The white curve represents the predicted average curved and the ribbons indicate a range of credibility intervals (CI) to show the variation around this mean. The dashed line indicates the location of the peak ($D^*$) and the shaded region defines the parameter space between $d_1$ and $d_2$ (see Online Resource 2, Analytic Results).

### S3 Analytic Results

In many contexts, a quadratic function is suitable for modeling disturbance curves without the use of a logistic link function. In this setting, we argue that a concave disturbance function may support the Intermediate Disturbance Hypothesis (IDH). Starting with the quadratic model, we solve for the disturbance $D$ when the slope is zero

$$\beta_D + 2\beta_{D^2} D = 0. \tag{36}$$

Solving for $D$, the critical point of the curve is

$$D_0 = -\frac{\beta_D}{2\beta_{D^2}}. \tag{37}$$

To observe a concave curve, the following conditions must hold

1. **Concavity Criterion**: $\beta_{D^2} < 0$, ensuring that the quadratic function opens downward.
2. **Range for $D^\star$**: The critical point $D^\star$ must lie when the disturbance bounds $(d_1, d_2)$, defined by

$$d_1 < D^\star < d_2. \tag{38}$$

Substituting the solution above, this condition becomes

$$d_1 < -\frac{\beta_D}{2\beta_{D^2}} < d_2. \tag{39}$$

This implies the following constraints on $\beta_D$

$$\beta_D > (-2\beta_{D^2})d_1 \tag{40}$$

$$\beta_D < (-2\beta_{D^2})d_2. \tag{41}$$

To satisfy the IDH requirements, all criteria must hold simultaneously

$$\begin{cases} \beta_{D^2} < 0, \\ \beta_D \in (-2\beta_{D^2}) \cdot (d_1, d_2) \end{cases} \tag{42}$$

This framework identifies the parameter space for $\beta_D$ and $\beta_{D^2}$ that supports an IDH curve. The bounds $(d_1, d_2)$ represent the disturbance levels within which the peak of the convex curve occurs.

### S3.1 Eliminating Concavity

Recall that the model is

$$\mu = \phi(\gamma) = \frac{K}{1 + \exp(-\gamma)} \tag{43}$$

where

$$\gamma = L(D) = \alpha + \beta_D D + \beta_{D^2} D^2. \tag{44}$$

Applying the logistic function yields disturbance curves that cannot by definition be concave. Thus a more general criterion is needed to determine the presence of an IDH curve.

**Definition** (unimodality). A function $f(D)$ is called unimodal on $[0,1]$ if for some value of $D^* \in [0,1]$, it monotonically increases for $x < D^\star$ and monotonically decreases for $x < D^\star$.

**Definition** (IDH function). A disturbance function $\mu = \psi(D)$ supports the IDH if it is a unimodal function on $[0,1]$ that reaches its maximum at $D^\star \in (d_1, d_2)$, where $0 \leq d_1 < d_2 \leq 1$.

One needs to examine monotonic properties of the function $\phi(L(D))$ defined by equations 43 and 44. Since $\phi(\cdot)$ is a monotonically increasing function on $R$, monotonic properties of the function $\phi(L(D))$ are determined by monotonic properties of the function $L(D)$, which is quadratic. As a result, a disturbance function 43 - 44 supports the IDH if two conditions hold

- The quadratic function $L(D)$ opens downward, which entails

$$\beta_{D^2} < 0 \tag{45}$$

- The point of maximum $D^\star$ lies inside the interval $(d_1, d_2)$.

Since $D^* = -\frac{\beta_D}{2\beta_{D^2}}$, the latter condition gives

$$\beta_D \in (-2\beta_{D^2} d_1, -2\beta_{D^2} d_2) \tag{46}$$

Overall, conditions 45 and 46 identify the parameter region that supports the IDH.
